# An archaic reference-free method to jointly infer Neanderthal and Denisovan introgressed segments in modern human genomes

**DOI:** 10.1101/2025.03.17.643330

**Authors:** Léo Planche, Anna Ilina, María C. Ávila-Arcos, Flora Jay, Emilia Huerta-Sanchez, Vladimir Shchur

## Abstract

Admixture between populations is a common feature of human history. Admixture events introduce new genetic variation that can fuel evolution. Characterizing the significance of admixture events on the evolution of a population across various species is of great interest to evolutionary geneticists. Local Ancestry Inference (LAI) methods infer genetic ancestry of an individual at a particular chromosomal location. Certain methods specialize in detecting archaic introgression, which consists of interbreeding between modern and archaic humans like Neanderthals and Denisovans. Most current LAI methods allow the detection of a single archaic ancestry, and post-processing may distinguish between multiple waves of introgression. These methods vary in how they choose archaic or modern reference genomes for the inference. Here, we present a new HMM-based method (DAIseg), which has the advantage of simultaneously distinguishing between multiple waves of ancient and recent admixture, using only modern human reference genomes. Simulations demonstrate that DAIseg achieves higher overall performance than state-of-the-art methods. We also apply DAIseg to Papuan populations to jointly detect Denisovan and Neanderthal introgressed segments, and identify a higher number of archaic segments than previous methods. Analysis of inferred introgressed segments, shows that we can identify evidence for two Denisovan introgression events in Papuans without having any post-processing and filtering. Overall, on top of being able to deal with both Archaic and recent admixture, DAIseg provides a more principled approach for detecting and classifying Denisovan and Neanderthal segments which will improve downstream analysis of introgressed segments to infer the impact of archaic introgression in humans.

## 1 Introduction

Throughout human history, admixture—the exchange of genes— has played a crucial role in shaping the genetic diversity observed in modern human populations [1, 2, 3, 4]. For instance, Neanderthals—identified through archaeological findings—and Denisovans—discovered through sequencing a genome from a small finger bone—diverged from the lineage of anatomically modern humans (AMH) approximately 450,000 to 800,000 years ago, with varying estimates across studies [5, 6, 7]. Thanks to advances in genomic sequencing, we now have genetic material from both of these extinct archaic populations [8, 9, 10, 11], enabling us to investigate their genetic relationships with AMH. Analyses of these ancient samples, alongside data from modern human populations, reveal that individuals of non-African origin carry approximately 2–3% Neanderthal DNA. Recent studies also report a lower prevalence (0.2%) of Neanderthal DNA in some African populations [12]. Denisovan genetic ancestry is predominantly found in populations from Oceania, South and Eastern parts of Asia, and the Americas [13, 14].

Quantifying and studying archaic introgression can be approached in two main ways. The first, global ancestry inference, estimates the proportion of an individual’s genome derived from specific ancestries, such as Neanderthal or Denisovan. The second, local ancestry inference (LAI), focuses on identifying the specific genomic regions inherited from these archaic populations which is the focus of this paper. Knowing where in the genome individuals carry archaic segments is helpful for providing critical insights into the role of natural selection acting on introgressed fragments (adaptive introgression), the timing of introgression events, the identification of introgression deserts in modern human genomes, the number of archaic sources and estimates of genetic diversity in archaic populations based on surviving fragments in modern genomes. Since only a limited number of archaic genomes have been sequenced, these fragments offer a unique opportunity to infer genetic diversity in archaic populations that interbred with AMH.

In LAI methods, chromosomes are conceptualized as mosaics of genomic tracts originating from multiple ancestral groups [15, 16]. LAI is a valuable tool for understanding population evolution, migration history, and disease risk [17]. Many LAI models rely mostly on Hidden Markov Models (HMMs), where hidden states represent ancestries and emissions correspond to observed haplotypes/genotypes [18, 19, 20, 21]. The ancient divergence between AMH and archaic populations results in substantial genetic distance, facilitating the detection of archaic ancestry in modern humans. However, the limited number of available archaic reference genomes may not fully capture the diversity of archaic individuals who actually contributed to modern humans, potentially underestimating archaic ancestry [9]. Despite these challenges, various methods have been developed to detect archaic fragments in modern genomes using archaic reference genomes [22], [23, 24, 25]. Alternative approaches, such as HMMix, identify archaic ancestry without using Neanderthal or Denisovan genomes, instead using a modern human population without introgression as an outgroup (e.g., African populations) [25]. While HMMix enables the detection of archaic fragments, its application often requires multiple runs and/or extensive post-processing to distinguish Neanderthal from Denisovan ancestry, making comparisons across studies challenging [26]. Classifying whether archaic segments come from Neanderthals or Denisovans is important because these segments are used to infer the time of introgression, the number of introgression events, and the genetic relationships between archaic populations.

Different studies have used slightly different approaches for classifying archaic segments. For example, to identify Denisovan ancestry in Papuans, [25] first ran HMMix to identify archaic segments with Africans as an outgroup, followed by another run using Africans, Europeans, and East Asians as outgroups. The segments that are detected only in the first run are classified as Neanderthal, and the segments detected in both runs are classified as Denisovan. Since, the second run integrates Eurasians into the outgroup, it effectively identifies archaic tracts that are in Papuans and absent in Eurasia which are assumed to be from Denisovans. In a recent study, [27] also applied HHMix to Papaun genomes, and they ran HMMix once followed by additional filterings to distinguish between Neanderthal and Denisovan segments. This led to some unexpected results; despite Papuans harboring more Denisovan ancestry, the majority of archaic segments identified are genetically closer to the Neanderthal genome than the Denisovan genome.

Another difference between studies is whether Papuans have experienced two introgression events from distinct Denisovan populations. For example, [28] found evidence for two introgression events, but Browning et al. does not identify evidence for more than one Denisovan introgression. There are several reasons why studies have different results. It may be due to differences in postprocessing steps required to classify archaic segments into Denisovan or Neanderthal segments or the fact that Denisovan and Neanderthal ancestry are not jointly inferred. Here, we present a generalized model (DAIseg) that jointly detects multiple ancestries in a single analysis, which enables the simultaneous identification of Neanderthal and Denisovan ancestry. We also show how this model provides a framework to study archaic ancestry in recently admixed genomes.

In this study, we benchmark our method using simulated data to evaluate its power to detect introgression and compare it to HMMix. We also apply our model to Papuan genomes—known to harbor both Neanderthal and Denisovan ancestry—and demonstrate that it can infer these ancestries simultaneously. Moreover, our method identifies additional introgressed segments not identified by HMMix. Our analysis of introgressed segments in Papuan genomes supports the hypothesis of two distinct Denisovan introgression events, consistent with previous findings. Additionally, the inferred Denisovan segments show greater genetic similarity to the sequenced Denisovan genome, as expected. Notably, these results are achieved through a single application of our method, reducing the need for the complex filtering and post-processing steps required by earlier approaches.

In summary, our approach offers a streamlined and powerful framework for jointly detecting and classifying archaic genomic segments as Denisovan or Neanderthal in origin. Applying this method to Papuan genomes, we identified more archaic fragments and uncovered clear evidence of two separate Denisovan introgression events. Finally, we show how our framework can be extended to deconvolute both archaic and recent modern human ancestry within recently admixed individuals. Our results highlight the utility of our method for advancing our understanding of the role of archaic ancestry in modern human evolution.

## 2 Model

### 2.1 General overview

In this subsection we provide intuition for the key ideas underlying the algorithm and in the next subsection 2.2 we describe the mathematics in more details. Our method generalizes HMMix [25] to enable the inference of archaic ancestry from multiple archaic populations. The main idea underlying HMMix is that archaic genomic segments introgressed into a population are expected to have a high density of private variants relatively to modern human populations which did not interbreed with Neanderthals or Denisovans. In HMMix, there are two hidden-states, Ingroup and Archaic. To construct observations, the genome is binned into windows of fixed length and the number of private variants is counted in each window. Emission probabilities are modelled with Poisson distribution parameterised by the mutation rate and the timing of admixture. Transitions between hidden states are caused by recombinations, following ancestral lineage switching its ancestry with the probability equal to the admixture proportion.

Our method generalises HMMix by allowing complex rooted population scenarios with multiple reference and founding populations as well as multiple admixture events. Target population is the population in which we are detecting introgressed segments for some of its individuals (similar to an ingroup in HMMix). Reference populations are populations used to construct observations by constructing sets of private variants of a target genome relatively to each of them (similar to an outgroup in HMMix). Ancestral populations define the set of hidden states, and they match the possible paths of ancestral lineage from target genome to the root of the population scenario. Reference populations can be either non-admixed outgroups as in HMMix as well as genomes representing ancestral populations (the benefit of including a small number of archaic genomes in the analysis is shown in [29]).

A population discriminates well between two ancestries if its average coalescence times with the two ancestral populations (taken separately) differs in a significant way. Our model distinguishes between *N* different ancestries using *M* reference populations. There is no necessary relationship between *N* and *M*. For example, it is possible to distinguish between more than two ancestries using a single reference population, or to use multiple reference populations to distinguish between only two ancestries, although in general *N* and *M* should be of the same order. In practice, when building the model, the reference populations should be chosen such that each discriminates well between at least two ancestries.

Let us now give a toy example to illustrate all of this. Assume a population such as admixed Americans, which received admixture from Africans, but which also has Neanderthal genomic segments from its Eurasian ancestry. Assume we are trying to infer Africans, Neanderthal and non admixed segments in that population. As there is little genetic data for Neanderthal, we will use modern Africans and modern Eurasians as reference populations. The target is admixed American, the ancestries are target, African, Neanderthal and the reference populations are African and Eurasian. We cut the genomes in small windows, and for each window, we remove on one side the variants present in Africans and on another side the variants present in Eurasia. Very roughly and intuitively, here is how the three ancestries will be detected: if a segment has very little variants that are absent in Africa, but more variants that are absent from Eurasia, then it is most likely inherited from Africa. If a segment has a large amount of variants absent in Africa, but little absent in Eurasia, then it is likely of Neanderthal origin. Finally if a segment has very little variants that are absent in Eurasia, but more variants that are absent from African, then it is most likely non introgressed. Such a demography is illustrated in figure 1b (one archaic and two reference populations). We also provide in figure 1a a demographic model, for the detection of Archaic ancestries in Papuans (two archaics and two reference populations). These two cases will be developed further in the following sections.

**Figure 1:**
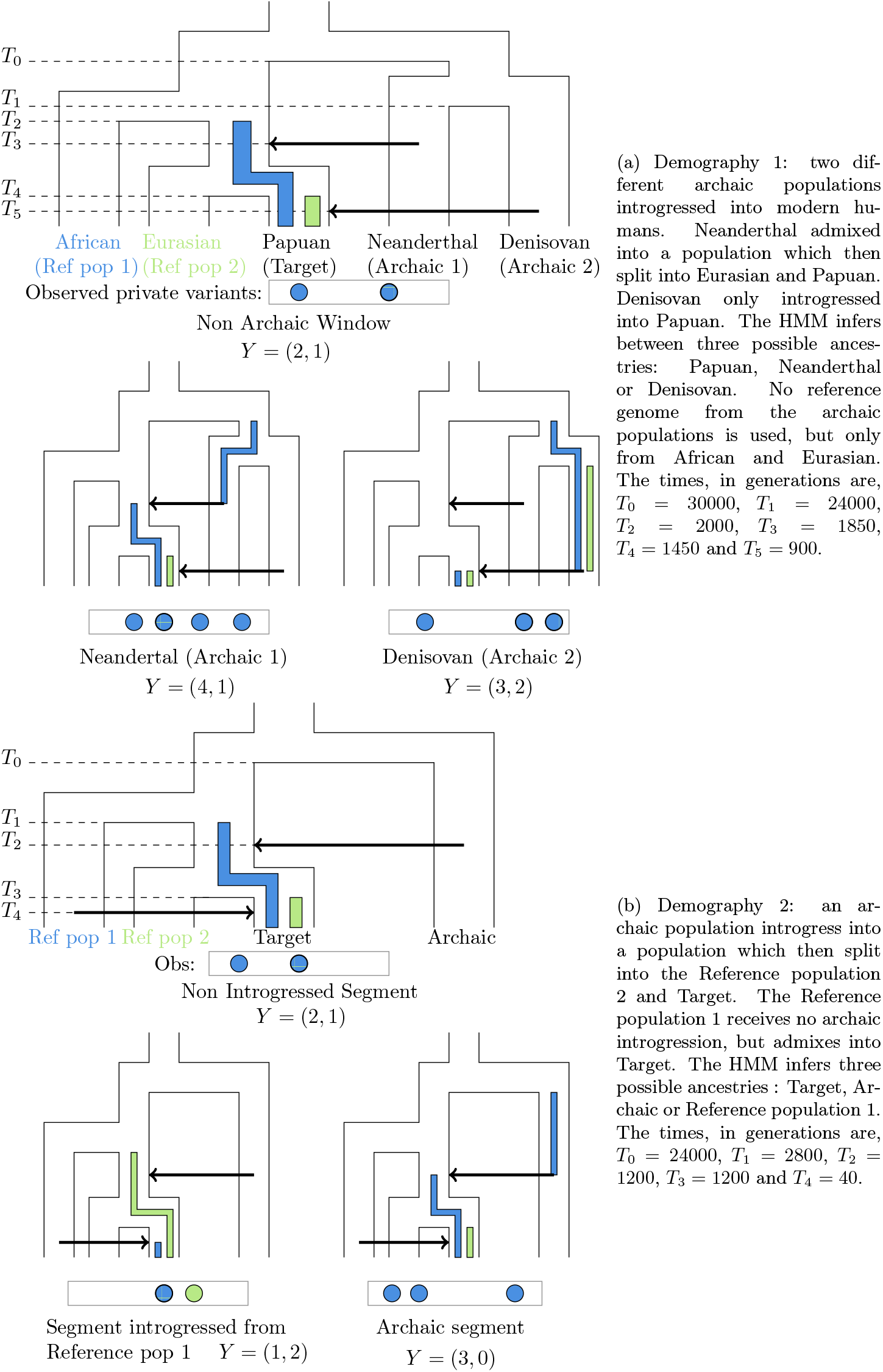
Observations used by the HMM. In both demographies, three possibles ancestries can be found in the Target. For each ancestry the blue and green shows the time of divergence between Target and the corresponding blue/green reference population. The higher the time of divergence the more private variants accumulate in the Target window, which is illustrated with the Observed private variants schema. The exact number of private variants with respect to each ancestry, *Y* = (*y*_1_, *y*_2_), is purely illustrative.

### 2.2 HMM architecture

In a similar fashion as in HMMix [25] the phased haplotype is binned in windows of constant size *L* and for each window we separately count the number of private derived variants in the target haplotype that are absent from each of the *M* reference populations, each represented by a certain number of individuals. These values are binned into a *M* -tuple, noted *Y*, which is the observation associated to the window. Note that the numbers of private variants relatively to each reference population are not independent values. In this section, we will approximate them as independent, which already provides good accuracy and in Supplementary materials we will detail how to take into account the dependency under certain scenarios and highlight its slight improvement. We illustrate the model with two examples in figure 1

#### 2.2.1 Emission probabilities

For an ancestry *x*, the probability of observing *Y* = (*y*_1_, *y*_2_…*y*_*M*_) is computed by multiplying the probabilities of observing each *y*_*i*_ given that the ancestry of the window is *x*. Let *µ* denote the mutation rate and 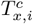 denote the mean coalescent time between ancestry *x* and reference population *o*_*i*_. The number *y*_*i*_ of private variants relative to population *o*_*i*_ follows a Poisson law with parameter 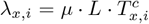. Let *Z*_*x,i*_ be a discrete random variable following a Poisson law with parameter *λ*_*x,i*_. We then get

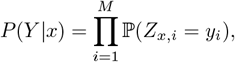

which concludes for our emission probabilities.

#### 2.2.2 Transition probabilities

For the transition probabilities, we expand the ideas from [25] to be able to deal with multiple admixture events. In this section, we use an example with two admixture events to explain the key ideas. The exact probabilities in the general case are given in Supplementary materials.

Consider the scenario with a target population *x*_0_ which received admixtures from populations *x*_1_ and *x*_2_. The first (oldest) admixture was with *x*_1_ at time *t*_1_ and with admixture proportion *a*_1_, the second, with *x*_2_, at time *t*_2_ and with proportion *a*_2_.

Let us compute the transition probability from ancestry *x*_1_ to *x*_0_. Such a transition is due to a recombination event which either took place between time 0 (present) and *t*_2_ or between time *t*_2_ and *t*_1_.

In the first case, the probability of recombination between 0 (present) and *t*_2_ is 1 − exp(−*rLt*_2_). We are only interested in recombination events into ancestry *x*_0_. Therefore the lineage should go toward the target population at both *t*_2_ and *t*_1_, this has probability respectively (1 − *a*_2_) and (1 − *a*_1_). Overall we get the probability of such an event of (1 − exp(−*rLt*_2_))(1 − *a*_1_)(1 − *a*_2_).

In the second case, the probability of recombination between *t*_2_ and *t*_1_ is exp(−*rLt*_2_)(1 − exp(−*rL*(*t*_1_ − *t*_2_))), where the first multiplier is the probability of no recombination before *t*_2_. We are already over the time point *t*_2_, so only admixture at *t*_1_ matters. As a result, we have the probability of such an event of exp(−*rLt*_2_)(1 − exp(−*rL*(*t*_1_ − *t*_2_)))(1 − *a*_1_).

Therefore, the final transition probability is (1 − exp(−*rLt*_2_))(1 − *a*_1_)(1 − *a*_2_) + exp(−*rLt*_2_)(1 − exp(−*rL*(*t*_1_ − *t*_2_)))(1 − *a*_1_).

## 3 Results

### 3.1 Simulations

We illustrate the flexibility and performance of our method by running simulations under two demographic scenarios shown in figure 1. The first scenario models two pulses of archaic introgression from two different archaic populations. The second scenario models one archaic and one modern admixture. Moreover, for the scenario 1b Reference population 2 is both an ancestry and a reference population.

#### 3.1.1 Demography 1

We test the method on the first demographic model (Figure 1a), it corresponds to the simple case of Papuan populations which have both Neanderthal and Denisovan ancestry in significant proportions. We wish to distinguish between three different ancestries, Neanderthal, Denisovian and anatomically modern human (AMH). To do so, as in [25], we will use two outgroups, Africans (Reference population 1) which received no archaic introgression and Europeans+East Asians (Reference population 2) which received introgession from Neanderthal (Archaic 1), but no or very little from Denisovan (Archaic 2). The African population allows us to discriminate AMH ancestry from archaic while the Eurasian population allows us to discriminate Denisovian ancestry from the rest. We simulate 150 individuals for each reference population and 20 individuals for the target population on 250Mbp and compare the performance of HMMix [25] and our model, DAIseg, on this dataset. The script details and availability can be found in (Supplementary materials).

As HMMix cannot natively distinguish between more than two ancestries, we run it twice, firstly to detect archaic segments, secondly to detect Denisovan segments. Afterwards, the segments that were classified as archaic, but not Denisovan are classified as Neanderthal. The results are shown in table 1.

**Table 1:** Comparison of DAIseg and HMMix methods for posterior detection of two components of archaic introgression (Neanderthal and Denisovan) based on 40 simulated haploid individuals (Demography 1). Precision, recall and F-score for Neanderthal, Denisovan and Archaic segments are given for each method with posterior threshold values of 0.5 and 0.98. Archaic segments are those classified as either Neanderthal or Denisovan. F-score is a summary statistic of Precision and Recall defined as 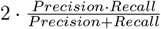.

|  |  | Posterior probability threshold |  |  |  |  |  |
| --- | --- | --- | --- | --- | --- | --- | --- |
|  |  | 0.5 |  |  | 0.98 |  |  |
|  |  | Precision | Recall | F-score | Precision | Recall | F-score |
| DAISeg | Neanderthal | 0.92 | 0.75 | 0.83 | 0.93 | 0.74 | 0.82 |
|  | Denisovan | 0.82 | 0.96 | 0.88 | 0.83 | 0.95 | 0.89 |
|  | Archaic | 0.96 | 0.996 | 0.98 | 0.98 | 0.99 | 0.98 |
| HMMix | Neanderthal | 0.94 | 0.62 | 0.75 | 0.96 | 0.64 | 0.77 |
|  | Denisovan | 0.75 | 0.99 | 0.85 | 0.78 | 0.98 | 0.87 |
|  | Archaic | 0.97 | 0.996 | 0.98 | 0.99 | 0.98 | 0.98 |

We show that combining all the information about outgroups / reference populations in a single model avoids extra post processing steps while slightly increasing the accuracy of the inference, as measured by the F-score. Note that when increasing the cutoff of the HMMs, i.e. the minimal outputted probability above which confidently assigning an introgressed ancestry, we expect an increase in Precision, but a decrease in Recall. This is true for all cases, but one, detecting Neanderthal segments using HMMix, where both precision and recall increase with the cutoff. Understanding this phenomena will make more intuitive the advantages of combining all reference populations in a single HMM. Indeeed, the most common reason why Neanderthal segments are underdetected in this particular demographic scenario is that they are mistaken for Denisovans (see confusion matrices in Supplementary materials). By increasing the cutoff in HMMix, one Neanderthal segment which was misclassified as Denisovan, might not be anymore, but still be detected as archaic. In other words this segment looked like Denisovan, but even more so Neanderthal. While our model will use this combined information to classify it as Neanderthal, two independent runs of HMMix will simply classify it as both archaic and Denisovan, hence Denisovan.

##### Unphased data

The method assumes that the data is phased, but we show, both with simulations and real data that it can be used on unphased data as well. When constructing the observations, we simply count all the private variants in the window. Under the current demography, the model still outputs three possible ancestries, Modern, Neanderthal or Denisovan while in fact any position in the genome has two ancestries (from each haplotype). In particular, if a position has both Denisovan and Neanderthal ancestries, the model will not be able to detect both ancestries and the recall score will inevitably decrease. It will still very likely be detected as Denisovan or Neanderthal, hence correctly identified as Archaic. We ran the model on simulated unphased data and the results are shown in table 2. Using unphased data mostly decreases the precision score for the archaic ancestries. This result can be intuitively understood. When using unphased data, we are summing the private variant present on both copies of the chromosome, increasing the number of observations per window. On this particular demographic model, a large number of observation per window is a sign of archaic ancestry.

**Table 2:** Precision and recall scores for DAIseg applied to phased and unphased simulated data. 20 Papuan-like diploid individuals were simulated using msprime [30] with two introgressed archaic populations corresponding to Neanderthals and Denisovans.

|  | Phased |  | Unphased |  |
| --- | --- | --- | --- | --- |
|  | Precision | Recall | Precision | Recall |
| Neanderthal | 0.92 | 0.75 | 0.91 | 0.73 |
| Denisovan | 0.82 | 0.96 | 0.77 | 0.95 |
| Archaic | 0.96 | 0.996 | 0.94 | 0.997 |

#### 3.1.2 Demography 2

We now test the model on the second demographic model. In this case the target population received both an archaic admixture and a modern one. The Archaic population split from modern humans 24000 generations ago and admixed 1200 generations ago. The Reference population 1 split from the target population 2800 generations ago and admixed 40 generations ago. In figure 1b we propose two reference populations to distinguish between the three possible ancestries in the target population. One can note that Reference population 1 theoretically contains already enough information to infer between the three ancestries. But due to the stochastic nature of the process, which can for example lead to incomplete lineage sorting, it is better to use as much relevant data as possible. Using Reference population 2 offers extra information which reduces uncertainty. This illustrates our previous point when presenting the HMM model, there is no direct relationship between the number of reference populations and the number of ancestries, our model can take into account any relevant information available. To illustrate this, we first run DAIseg using only Reference population 1 (the observations are then one integer per window) and then using both Reference populations 1 and 2 (the observations are then two integers per window), all other parameters being equal. The results are summarized in Table 3. Notice that using only one reference population our method already achieves high accuracy for distinguishing between three ancestries. Adding a second reference population does indeed increase the overall inference accuracy. At the same time this additional Reference population 2 does not help to increase the accuracy for the detection of Archaic segments (precision of 0.94 and recall of 0.97 in both cases). The reason being that Reference population 1 already has a very large divergence time with that population, yielding a high number of private variants. On the other hand, using Reference population 2 helps to distinguish between Target and Reference population 1 segments.

**Table 3:** Precision and recall scores for DAIseg applied to the simulated data under Demography 2. In the left column only reference population 1 was used to construct observations. In the right column reference populations 1 and 2 were jointly used for the inference which helps to better distinguish between Target and Reference population 1 ancestry.

|  | Reference populations used |  |  |  |
| --- | --- | --- | --- | --- |
|  | Reference pop 1 |  | Reference pop 1 & Reference pop 2 |  |
|  | Precision | Recall | Precision | Recall |
| Target | 0.88 | 0.99 | 0.97 | 0.99 |
| Archaic | 0.94 | 0.97 | 0.94 | 0.97 |
| Reference pop 1 | 0.997 | 0.8 | 0.998 | 0.95 |

### 3.2 Application to Papuan genome

We apply our model to Papuans individuals — who harbor Denisovan and Neanderthal ancestry — from the Simons Genome Diversity Project [31] to detect and classify introgressed segments as being of Denisovan or Neanderthal origin in real genomic data. We use two outgroup populations, the first consists of 205 Africans from 1000 Genomes Project [32], from Yoruban (YRI) and Esan (ESN) populations. The second ougroup population consists of 814 Eurasians from British (GBR), Finnish (FIN), Toscan (TSI), Utah (CEPH) with Northern and Western European ancestry populations, Japanese (JPT) and Han Chinese (CHB). Our HMM has three hidden states, Modern, Neanderthal and Denisovan. Our underlying assumptions in choosing our outgroup populations are that African populations do not carry Neanderthal or Denisovan ancestry and that African+Eurasian populations do not harbor Denisovan ancestry.

We compare our model to HMMix which was ran twice; the first time using Africans as an outgroup, and the second time using Africans and Eurasians as the outgroup. We followed [25] procedure for classying archaic segments into Denisovan or Neanderthal: archaic segments detected only in the first run are classified as Neanderthal and archaic segments detected in both runs are classified as Denisovan. Using DAIseg, we detect 24% more Neanderthal segments and 17% more Denisovan segments than HMMix. Out of all Neanderthal segments detected with HMMix, we classify 82% of them as Neanderthal, 5% as Denisovan and 13% as non Archaic. For Denisovan segments detected with HMMix, we classify 93% of them as Denisovan, 1% as Neanderthal and 6% as non Archaic. To check if those differences are the result of miss-classification, we compare the similarity to archaic reference genomes of segments detected only by DAIseg, only by HMMix or by both methods. Here, for each segment, the similarity is computed by counting the number of SNPs in the segment where an allele is shared between the target Papuan and the Archaic (either Vindija Neanderthal or Altai Denisovan), then dividing by the total number of SNPs present in the segment. Overall, our model detects more archaic segments and closer inspection (Figure 2) shows that these are mostly of Archaic origin and not the result of misclassification. Figure 2 also shows that Neanderthal segments detected with DAIseg have slightly higher genetic similarity to the Neanderthal genome than Neanderthal segments detected with HMMix. For Denisovan segments identified with DAIseg, genetic similarity to the Denisovan genome is slightly lower than for Denisovan segments detected with HMMix.

**Figure 2:**
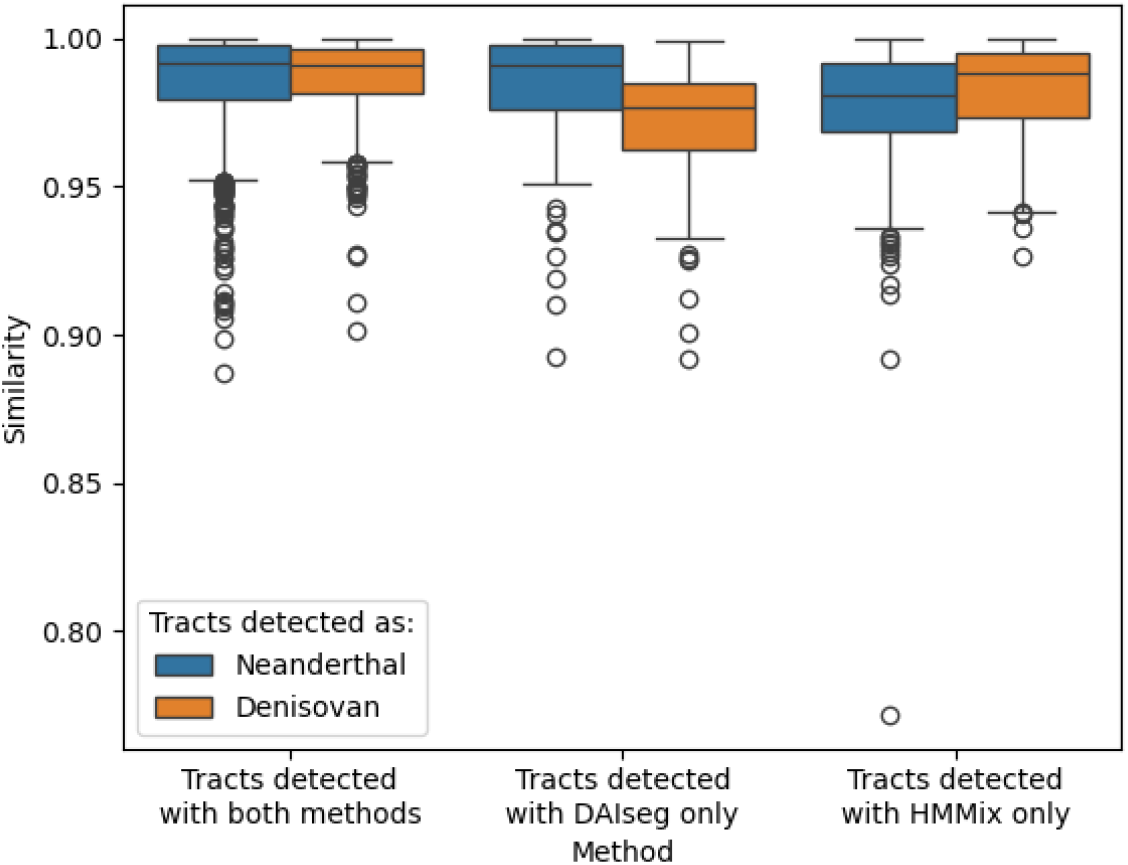
Similarity of the segments detected as Neanderthal (resp. Denisovan) to the Vindija Neanderthal (resp. Altai Denisovan) reference genome. On the left, the similarity computed for segments that are detected with both DAIseg and HMMix, in the middle the segments that are only detected with DAIseg, on the right, the segments only detected with HMMix. Segments detected with both methods account for 66% of all segments, segments detected only with DAIseg for 26% and detected only with HMMix for 8%. Here, all SNPs present in the segments are used to compute the similarity, non Archaic segments have a similarity of around 0.94 to archaic reference genomes.

As previous studies have identified evidence for two Denisovan introgression events into Papuans [28, 27, 33], we analyzed Denisovan segments detected using DAIseg to determine if two subsets of Denisovan segments exhibit varying levels of genetic similarity to the sequenced Denisovan genome. In figure 3, we take all Denisovan segments detected with DAIseg and compute the genetic similarity to the Denisovan genome (y-axis) and the Vindija Neanderthal genome (x-axis). For this plot, genetic similarity is computed using archaic private variant only, the details are given in supplementary materials 4. This also offers another way to check the accuracy of our model. On average we expect Denisovan segments to appear above the diagonal, which means more similarity to the sequenced Denisovan genome than to the sequenced Neanderthal genome, and this is what we observe. For Neanderthal segments, we expect them to appear below the diagonal (see second panel of Figure 3). Following [28] to determine if there is evidence for two waves of Denisovan introgression, we kept all the segments detected as Archaic with length greater than 200kb and conditioned on observing at least 10 private variants (with respect to African populations) in the Archaic genome. For completeness, we also give the same figure without any length threshold on the segments (figure 5).

**Figure 3:**
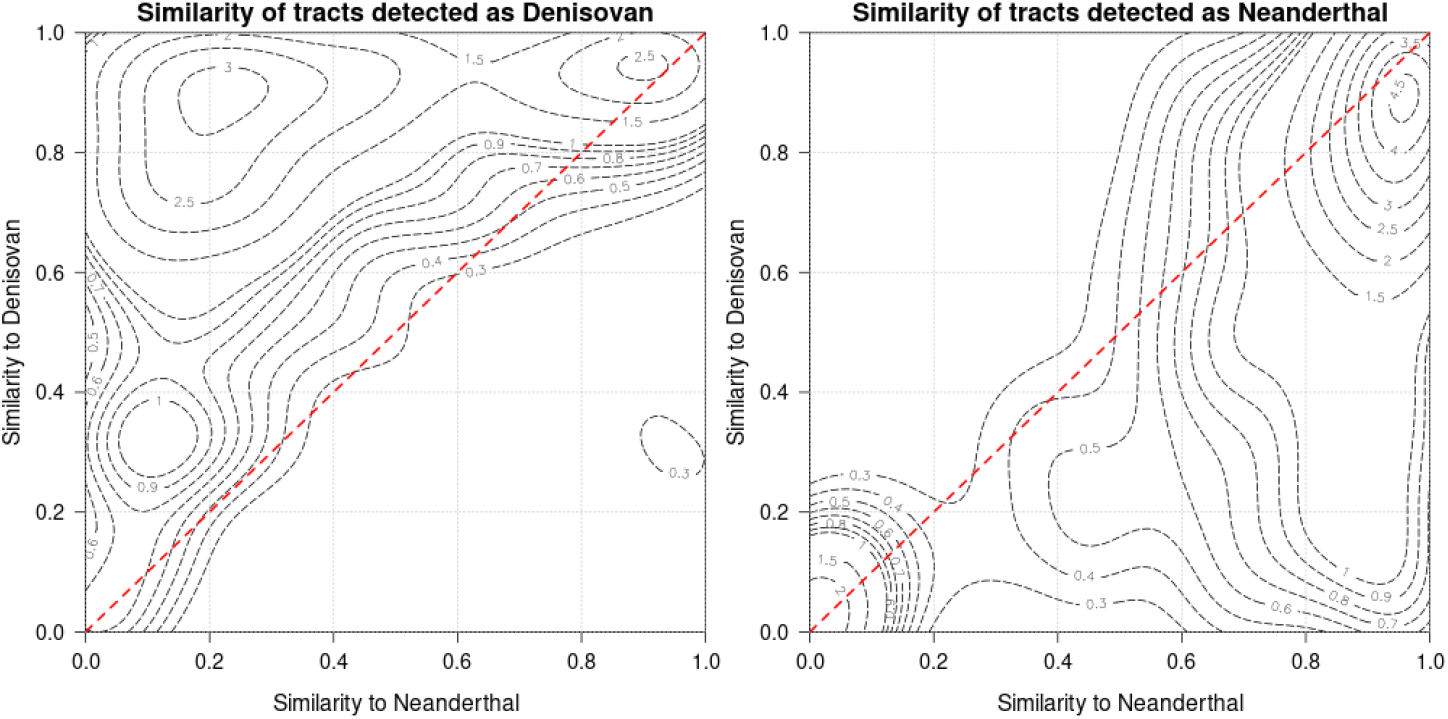
Similarity of the segments found in Papuan, using DAIseg, of size larger than 200kb, detected as Denisovan (left panel) or Neanderthal (right panel) to the Vindija Neanderthal (x-axis) and Denisovan (y-axis) reference genomes. 15% of segments detected as Neanderthal and 6% of segments detected as Denisovan shows a similarity lower than 0.2 to either Archaic genome. The red dashed line, *y* = *x*, correspond to an equal similarity to the Neanderthal and Denisovan reference genomes.

**Figure 4:**
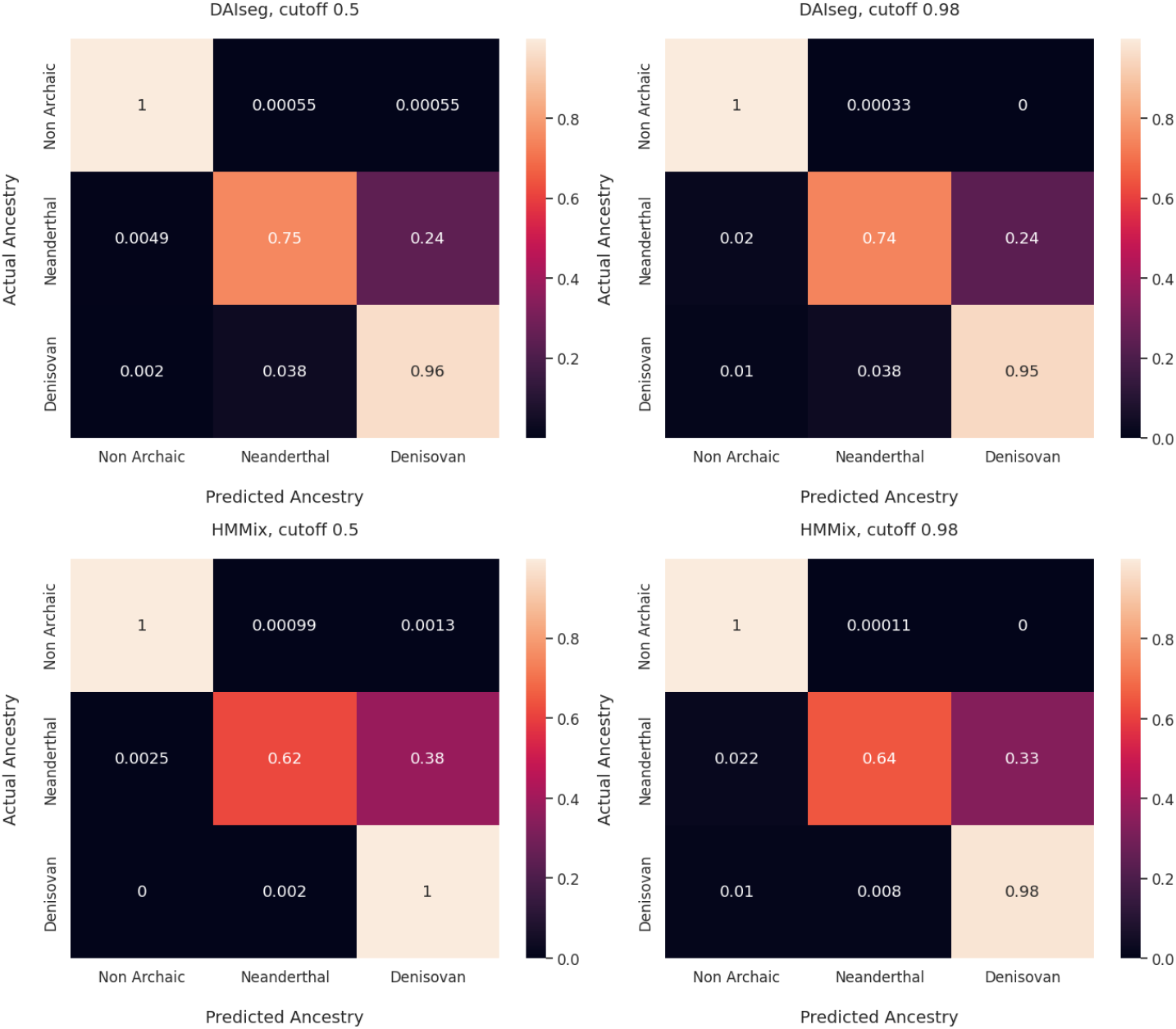
Confusion matrices for the ancestries inferred with DAIseg and HMMix based on simulations described in 3.1.1 with different posterior tresholds supporting archaic segments (0.5 and 0.98).

**Figure 5:**
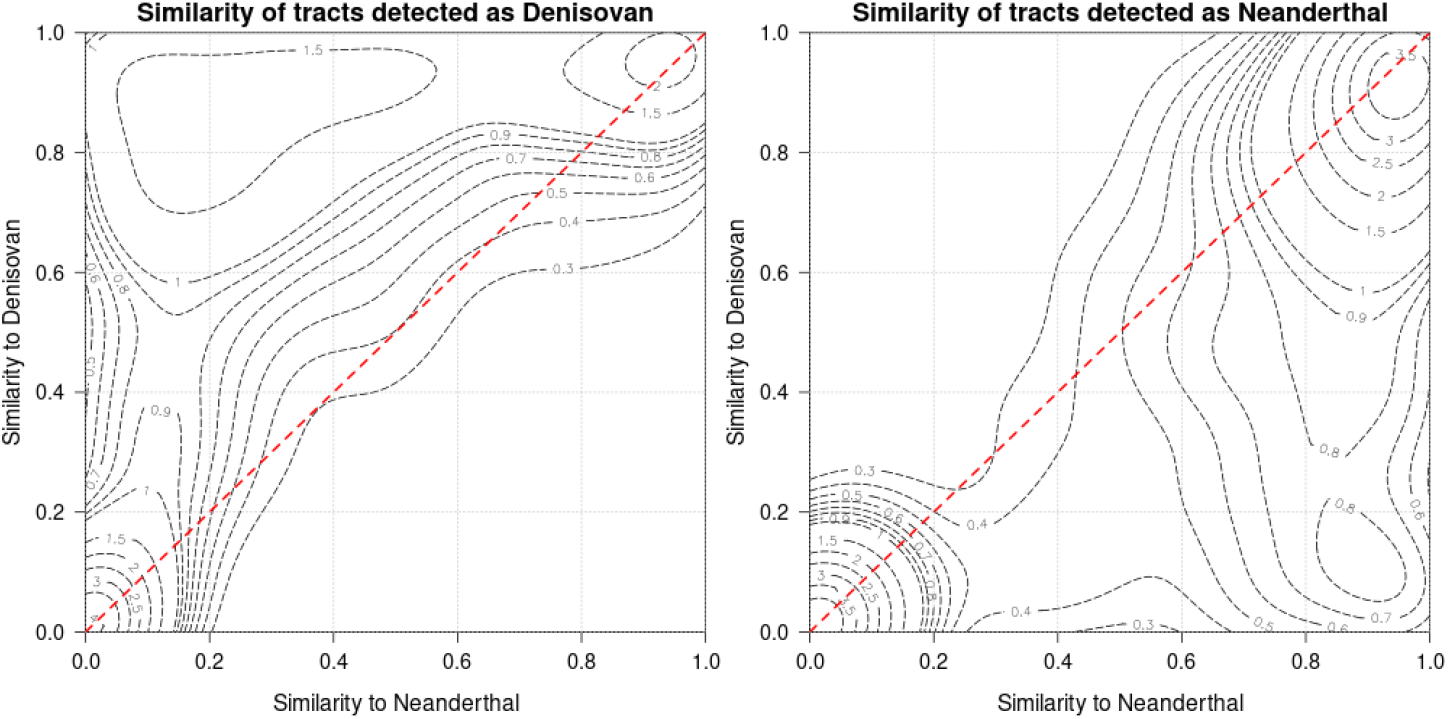
Similarity of the segments found in Papuan, using DAIseg, without any filtering on segment size detected as Denisovan (left panel) or Neanderthal (right panel) to the Vindija Neanderthal (x-axis) and Denisovan (y-axis) reference genomes. 96% of segments not detected as Archaic, 24% of segments detected as Neanderthal and 18% of segments detected as Denisovan shows a similarity lower than 0.2 to either Archaic genome. The red dashed line, *y* = *x*, correspond to an equal similarity to the Neanderthal and Denisovan reference genomes.

One can notice that for both Neanderthals and Denisovan there is slight peak in the distribution around the origin, which means low similarity to both of the reference sequences. By defining that peak as having similarity below 0.2 to both Archaic genomes, for segments detected as Neanderthal, around 15% are at this peak, for Denisovan segments around 6%, on the other hand, more than 97% for segments detected as non Archaic. This numbers can be interpreted as a rough estimate of the rate of false positive and false negative. The rate of false positive is higher for smaller segments, but even then, it does not exceed 18% for Denisovan segments and 24% for Neanderthal segments (figure 5).

Apart from the peak around the origin, for tracts detected as Denisovan, we observe three peaks, one near (0.9,0.9), one around (0.2,0.9) and one around (0.1,0.3), with the first number in the parenthesis being the similarity to Vindija Neanderthal and the second, the similarity to Altai Denisovan. For tracts detected as Neanderthal, no such behavior is observed, the similarity to Neanderthal show only one peak, at around 0.9. These observed varying levels of genetic affinity between inferred introgressed segments and sequenced archaic genomes is consistent with more than one introgression event with Denisovans and one introgression event with Neanderthals.

We compare the length (Figure 6 of the Neanderthal and Denisovan segments. We find the Neanderthals segments to be slightly longer than the Denisovans. The introgression map is available in Supplementary materials.

**Figure 6:**
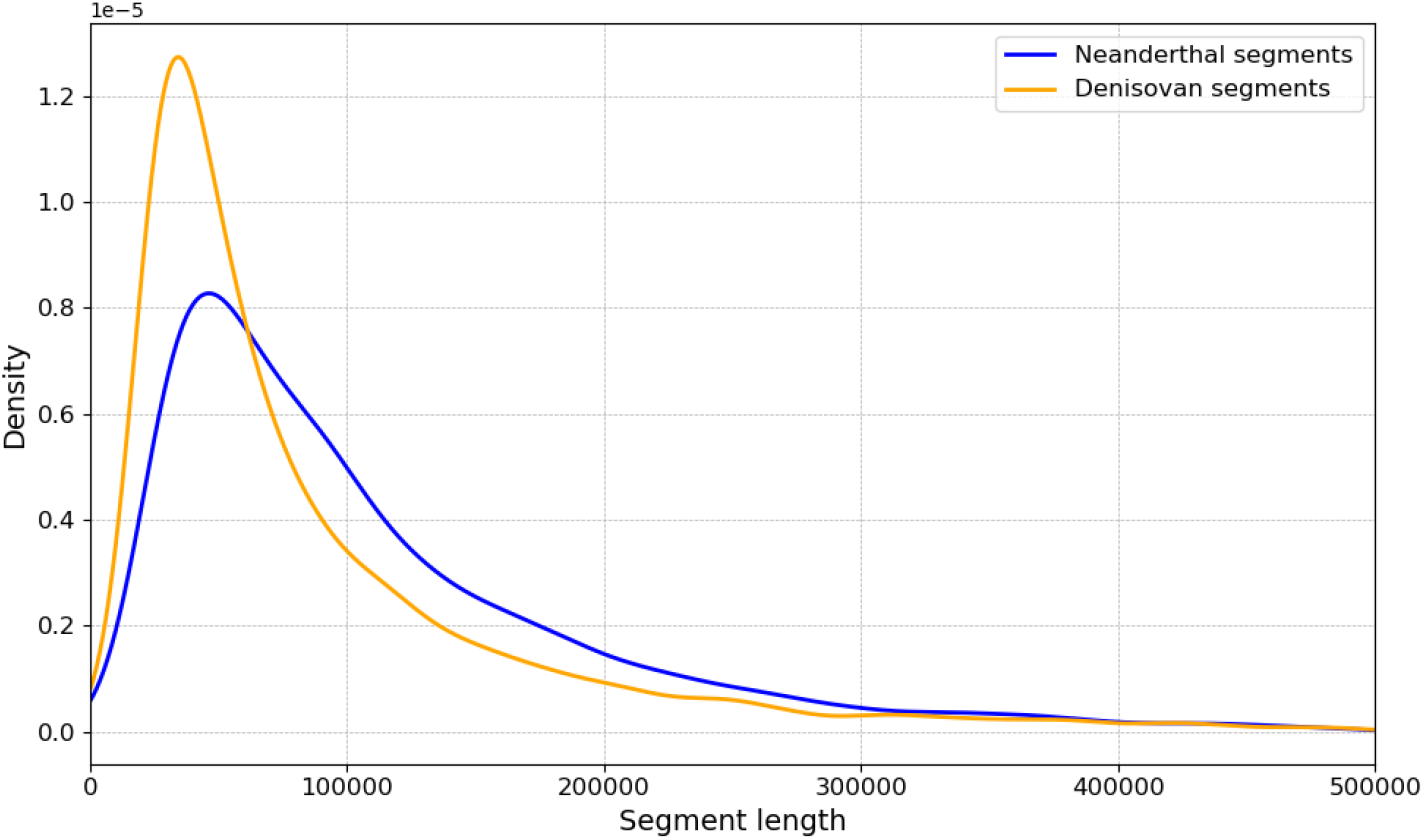
Denisovan segments are slightly shorter than Neanderthal segments. Average segment length is 109kb for Neanderthal and 87kb for Denisovan.

## 4 Discussion

Here we present a new method, DAIseg, to detect introgressed segments from multiple archaic sources. Our framework is general and can be extended to detect introgression from several sources, including non archaic ones. In the case of Neanderthal and Denisovan introgression (see Demographic model in Figure 1a), and using simulated data, we showed that our performance is similar or better than state of the art methods. For example, compared to HMMix, we get a similar F-score for Denisovan (0.88 vs 0.85) and Archaic detection (0.99 vs 0.99), but our performance is higher for Neanderthal detection (0.83 vs 0.75). One big advantage of our method is that DAIseg allows for the detection and classification of multiple ancestries with a single run of the HMM. While we mostly focused on Denisovan and Neanderthal introgression, we also showed how one can detect archaic ancestry in recently admixed individuals (Demographic model 1b). Using simulations, we showed that we can accurately detect African, European and archaic ancestry, suggesting that this framework can be used to deconvolute both archaic and modern human ancestry in recently admixed individuals.

One big advantage of our method is that it not only detects archaic segments but it also classifies segments as being from a Denisovan or Neanderthal source. This is particularly useful when the target individuals harbor ancestries from both of these archaic sources. Here, we applied DAIseg to genomic data from Papuans who are known to carry ancestry from both Denisovans and Neanderthals. We find that our method identifies around 18% more introgressed segments than HHMmix, likely because we are using more information when we jointly infer tracts. We infer the majority of these additional segments to be of Neanderthal origin, which we further confirmed by looking at the similarity of these tracts to Archaic reference genomes (Figure 2).

We note that around 25% of all inferred archaic segments have a very high similarity to both Neanderthal and Denisovan reference genomes. These segments are sometimes classified as Neanderthal and sometimes as Denisovan by our model (but always as Archaic), showing that these may be shared segments or they might have a more complex demographic history. Importantly, the majority of archaic segments detected by DAIseg align with expected features. For example, Papuans harbor more Denisovan ancestry (3.6% of their genomes) than Neanderthal ancestry (2.8%), which is consistent with prior findings. Moreover, inferred Denisovan segments are genetically closer to the sequenced Denisovan genome than to the Altai Neanderthal genome, while inferred Neanderthal segments show greater similarity to the sequenced Neanderthal genome (Figure 3). These observations confirm that DAIseg reliably classifies introgressed segments and provides a principled framework for segment detection without relying on extensive post-processing steps.

We note that a previous study [27] presented unexpected results when detecting archaic ancestry in Papuans. They identified 60% of archaic segments in Papuans as being more similar to the Neanderthal genome, even though reported estimates of Denisovan ancestry in Papuans are higher than Neanderthal ancestry. However, many of the segments they detect exhibit only slightly lower similarity to the Denisovan genome, which may correspond to segments that we infer as Denisovan but also have a high genetic similarity to Neanderthals (top right peak in Figure 3). By jointly inferring Neanderthal and Denisovan ancestry, DAIseg resolves such ambiguities and produces results that align with evolutionary expectations, offering an improvement over prior approaches. Another example of this regards the number of Denisovan sources that contributed to ancestral Papuan populations.

In line with previous studies, our results provide clear evidence of two separate Denisovan introgression events, without requiring *post hoc* filtering. Specifically, when considering inferred Denisovan segments Figure 3 reveals three peaks. One peak contains segments of high genetic similarity only to the Denisovan genome, another peak is made up of segments with high genetic similarity to both the Neanderthal and Denisovan genome, and a third peak has intermediate genetic similarity to only the Denisovan genome. When considering segments detected as Neanderthal, we find that genetic similarity is closest to Neanderthal (see figures 3) although some segments also have high similarity to the Denisovan genome. These signatures suggests that at least three archaic contact events occurred in ancestral Papuan populations—two with Denisovans and one with Neanderthals. Notably, this additional Denisovan pulse was not detected using Sprime segments to construct the contour plot in [23]. Finally, regarding the peak comprising segments that are highly similar to both Neanderthals and Denisovans, we currently interpret this as the result of having introgressed fragments that are present in both archaic populations due to shared ancestry or admixture events between archaic populations [34]. In [27], they propose an alternative explanation for these archaic segments, where they suggest an introgression event from an archaic population that is closely related to both Denisovans and Neanderthals. However, as most analyses have harsh thresholds—often based on genetic similarity—these segments are often missed or excluded, and future analysis of all archaic genomes may reveal the history of these introgressed segments.

We acknowledge that our model requires carefully chosen outgroup populations and a welldefined demographic history. Although DAIseg does not explicitly use Denisovan or Neanderthal genomes for inference, the method can easily be extended to include them as reference populations, and it offers substantial flexibility to incorporate diverse demographic scenarios. For example, a recent study demonstrated that incorporating multiple Neanderthal genomes as references improves the detection of Neanderthal introgressed segments [29]. While only a single Denisovan genome is currently available, the sequencing of additional Denisovan genomes in the future could further enhance the detection and classification of Denisovan ancestry. One exciting direction is to use DAIseg on ancient genomes. This would require taking into account missing data, but as more ancient genomes are being imputed, using the genotype probabilities might be sufficient. In sum, the success of our method in identifying multiple Denisovan introgression events in Papuans suggests its broader applicability. Applying DAIseg to other populations, such as South Asians or East Asians, may uncover additional instances of Denisovan introgression events, providing further insights into archaic admixture.

## Acknowledgement

AI and VS were supported by the RSF grant 22-71-10056. This work was supported by the Human Frontier Science Project grant (number RGY0075/2019).

## Supplementary materials

### Transition probabilities

We now detail the intuition given in 2.2. Let *x* be any ancestry. If *x* is not the target ancestry, then let *T*_*n*_ be its admixture time in the target population and *a*_*x*_ its admixture proportion. If *x* is the target ancestry, then *T*_*n*_ is the starting time of that population and *a*_*x*_ is equal to one. For any time *t* lower than *T*_*n*_, we note *a*_*x*_(*t*) the proportion of ancestry *x* in the target population at time *t*. Let *T*_1_, *T*_2_, …*T*_*n*−1_ be the time of all admixture events in the target population which happened between current time *T*_0_ and *T*_*N*_. For *i* ∈ {1…*n* − 1}, let *a*_*i*_ be the admixture proportion associated with the admixture event at time *T*_*i*_. Then for any time *t*, such that *T*_*z*_ ≤ *t < T*_*z*+1_,

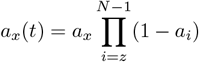

Let *x* and *x*^*′*^ be two possible ancestries in the model, if one of them is the target ancestry, then *T*_*x,x′*_ denotes the admixture time of the non target population into Target. If both *x* and *x*^*′*^ are admixed ancestries then *T*_*x,x′*_ is the time of admixture of whichever admixed last with Target.

With *r* the recombination rate, for a given window, the number of recombination event during time *T* is Poisson distributed with rate *r* · *L* · *T*. Therefore, the probability of having at least one recombination is 1 − *e*^−*r*·*L*·*T*^ which can be approximated by *r* · *L* · *T*. Let us define *T*_1_, *T*_2_, …*T*_*n*−1_ as before with *T*_*n*_ = *T*_*x,x′*_. The probability of having a recombination event from *x* to *x*^*′*^ is given by summing for each time interval [*T*_*i*_, *T*_*i*+1_] the probability of having a recombination event into *x*^*′*^ in that interval and no recombination afterward. This last probability being 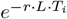 which can be approximated by 1 − *r* · *L* · *T*_*i*_. Finally, we get,

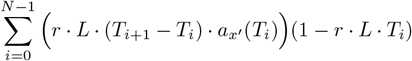

Which concludes for our transition probabilities.

### Conditional probabilities

In section 2.2 we defined the emission probabilities as if the *M* observations in a single window were independent values. Let us remember why it is not always the case. Let’s come back to the example of Papuans which split from Eurasians at time *T*_1_, while Eurasians split from Africans at time *T*_2_. It is clear that most of the private variants in Papuans relative to Eurasians are a subset of the private variants in Papuans relative to Africans. It follows that the number of private variants in Papuan relative to Eurasians and the number of private variants in Papuan relative to Africans are not independent values and can be computed using conditional probabilities.

We will explain in this section how to deal with condition probabilities in specific demographic models. This is the default behavior of DAIseg (when the demography is appropriate) and what we used for inference in real data. We will use conditional probabilities for demographies where one can sort the Reference populations such that after sorting, for each ancestry *x* and *i < j* the split time between reference population *i* and *x* is larger than the split time between reference population *j* and *x*. This is in particular the case for demography 1, corresponding to Papuan populations. Indeed, the African reference population has a larger split time with Papuan, Denisovan and Neanderthal compared to the Eurasian reference population.

When this condition is met, we sort in decreasing order the reference populations of our model in function of their split times as described above. The split time here being used as a first approximation or proxy for the mean coalescent time. For each window, we then compute the decreasing sequence *Y* = (*y*_1_, …, *y*_*M*_) such that *y*_*i*_ is the number of private variants found in the window relatively to the first *i* reference populations (after sorting). Let us note *µ* the mutation rate, 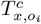 and the mean coalescent time between an ancestry *x* and reference population *o*_*i*_,

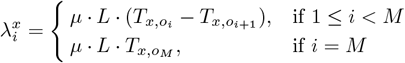

and

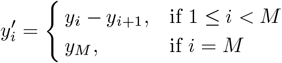

Where 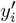 represents the number private variants which accumulated in our target population between times 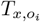 and 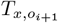. Finally, let *Z*_*i*_ be a discrete random variable following a Poisson law of parameter 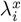, we then have:

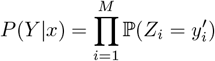

Which concludes for our emission probabilities. For demography 1 (fig. 1a), we applied DAIseg with emission probabilities defined as in the main text 2.2 and then with the conditional probabilities as defined here. The difference in accuracy is shown in table 4. Using conditional probabilities increases the F-score of the inference. By comparing these results to table 1, one can notice that running the model without conditional probabilities outputs scores extremely similar to HMMix. This should come to no surprise as results for HMMix are coming from running the HMM twice, independently.

**Table 4:**
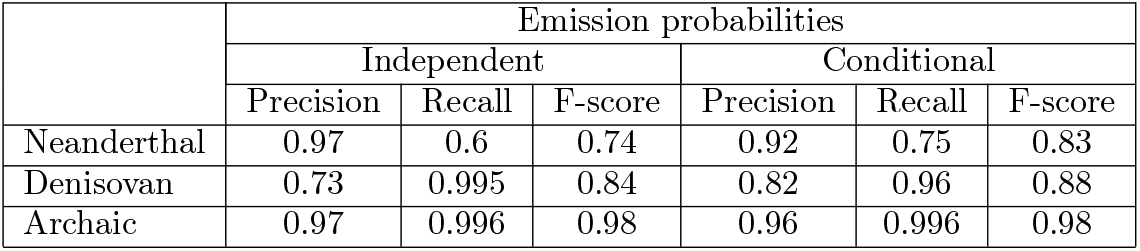
Precision and recall scores for DAIseg applied to the simulated data, using two different ways to define emission probabilities: independent (see section 2.2) and conditional (see section 4).

### Similarity to Archaic genomes

The similarity for a segment is computed as such: we keep all the positions with an archaic private variant i.e. a variant that is found in an Archaic genome, but absent in Africa. Then, there is a match if the archaic and Papuan individual share the private allele at this position. The similarity is the proportion of matches along all the positions containing an archaic private variant.

More precisely, let *m*_1_, …, *m*_*N*_ be modern African haplotypes, *a*_1_, *a*_2_ - archaic reference haplotypes and *p*_1_, *p*_2_ - Papuan haplotypes. Define the following sets of variable positions for a tract

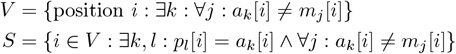

Then similarity is simply the ratio of the number of positions in each of the two sets

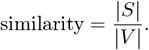

### Simulations

We used msprime [30] to simulate data. For the first demograhic model we simulated 20 Papuans, 150 Africans and 150 Europeans, using demographic parameters from [35] which are illustrated in figure 1a. For the second demographic model, we simulated 20 individuals from the Target population, 150 from Reference population 1 and 150 from Reference population 2. We took demographic parameters such that the Target population could represent native Americans who received admixture from African populations after the European colonization. The demographic parameters are illustrated in figure 1b. We use diploid genomes of size 250,000,000 bp, constant recombination rate *r* = 1.2 · 10^−8^ and mutation rate *µ* = 1.25 · 10^−8^. The simulation file is available at: https://github.com/LeoPlanche/DAIseg/tree/main/simulations/simulations.py

## Data availability

### Method

The method is implemented in Python and is available for use at: https://github.com/LeoPlanche/DAIseg

### Inferred segments

The inferred introgression map for archaic segments in Papuans is available at: https://github.com/LeoPlanche/DAIseg/blob/main/src/IntrogressionMap

The file will be gradually completed as more data becomes available.

## Supplementary figures

